# EuDockScore: euclidean graph neural networks for scoring protein-protein interfaces

**DOI:** 10.1101/2024.06.04.597410

**Authors:** Matthew McFee, Jisun Kim, Philip M. Kim

## Abstract

Protein-protein interactions are essential for a variety of biological phenomena including mediating biochemical reactions, cell signaling, and the immune response. Proteins seek to form interfaces which reduce overall system energy. Although determination of single polypeptide chain protein structures has been revolutionized by deep learning techniques, complex prediction has still not been perfected. Additionally, experimentally determining structures is incredibly resource and time expensive. An alternative is the technique of computational docking, which takes the solved individual structures of proteins to produce candidate interfaces (decoys). Decoys are then scored using a mathematical function that predicts the energy of the system, know as a scoring functions. Beyond docking, scoring functions are a critical component of assessing structures produced by many protein generative models. Scoring models are also used as a final filtering in many generative deep learning models including those that generate antibody binders, and those which perform docking. In this work we present improved scoring functions for protein-protein interactions which utilizes cutting-edge euclidean graph neural network architectures, to assess protein-protein interfaces. These euclidean docking score models are known as EuDockScore, and EuDockScore-Ab with the latter being antibody-antigen dock specific. Finally, we provided EuDockScore-AFM a model trained on antibody-antigen outputs from AlphaFold-Multimer which proves useful in re-ranking large numbers of AlphaFold-Multimer outputs. The code for these models is available at https://gitlab.com/mcfeemat/eudockscore.

## 1. Introduction

Protein-protein interactions are essential to a variety of biological processes including mediating chemical reactions, signaling, the immune response, and controlling the cell life cycle. Aberrations in protein-protein interactions are associated with a variety of diseases including cancers (L. Garner and D. Janda, 2011). To understand these interactions, the structures of individual proteins as well as protein complexes need to be obtained and studied. Historically, these structures were solved by using techniques such as X-ray crystallography, nuclear magnetic resonance, or cryogenic electronic microscopy. However, these techniques are expensive, time-consuming, and challenging leading to the development of a variety of computational techniques for structure prediction (Zhang, 2008).

### 1.1. An introduction to protein-protein docking and scoring

Protein-protein docking encompasses the development of computational pipelines to propose potential interfaced complexes of two or more solved individual protein structures, most often from the Protein Data Bank (PDB) (Burley *et al*., 2018). The docking process is typically broken down into two stages. The docking procedure proposes complexed structural models known as decoys, and the scoring stage (Tsuchiya *et al*., 2022). The scoring stage involves a physical model taking in structural variables (e.g. coordinates and atom types) and assigning a score to the complex indicating energetic favourability. Typically, to align with the concept of energies in physics, lower scores indicate more energetically favorable systems. These scoring functions are used to rank proposed models for selection and further investigation (Guedes *et al*., 2021).

### 1.2. An overview of scoring functions

In the past, scoring functions were typically hand-engineered to capitulate the known physics driving the folding and interactions of proteins. For illustrative purposes, a popular traditional scoring function is the Rosetta energy function (Alford *et al*., 2017), which includes terms involving the electrostatics of charged atoms, bond lengths, Van der Waals forces, etc. These models involved some model parameters being fit to available physical datasets (typically small). Although these models are useful, they often fail to capture the correct energetics of protein systems and fail to fully leverage the increasing amount of available biological data. Deep learning allows for the development of highly complex non-linear functions using available data for model parameter tuning.

Many earlier efforts for protein-protein interface scoring with deep learning involved discretization of protein-protein interfaces into 3D image-like grids to be input into convolutional neural networks (Schneider *et al*., 2021; Renaud *et al*., 2021; Wang *et al*., 2019). Due to the limitations of these models, researchers moved towards using graph neural networks (GNNs) which utilize mathematical graph representations of proteins as model inputs (Wang *et al*., 2021; Réau *et al*., 2022).

Continued efforts have been made to develop advanced graph neural network architectures to operate on graph representations at both the atom and residue levels for proteins. Modern architectures typically depend on equivariant mathematical operations derived from physics (Zitnick *et al*., 2022; Liao and Smidt, 2023; Liao *et al*., 2023) which can operate directly on input 3D coordinates without any necessary engineering. Typically these models will only need atom types and 3D coordinates to make predictions. Here equivariance means that any outputs of the model that are vectors will rotate appropriately with the inputs, furthermore, scalar outputs will be invariant.

### 1.3. Scoring functions in the era of AlphaFold

Generative deep learning models now dominate over traditional docking methods. For example, the bioinformatics field was revolutionized by the publication of AlphaFold2 (Jumper *et al*., 2021), a deep learning model which can take in protein sequences and produce highly accurate structures. The AlphaFold2 architecture was then extended to generate the structures of multi-chain protein complexes in the form of AlphaFold-Multimer (AFM) (Evans *et al*., 2022), providing an alternative avenue to computational docking techniques. Although AFM can produce accurate complex models in a large number of cases, there are many inputs in which it still struggles to correctly determine the interface (Evans *et al*., 2022) and quality metrics (iPTM, plDDT, etc.) are still necessary to differentiate high-quality from low-quality models. Another example is AbDiffuser (Martinkus *et al*., 2024), a deep learning model that generates antibody structures. It uses additional deep learning models that provide information such as predicted naturalness of predictions and predicted stability. Scoring models also remain a critical component of deep learning-based docking methods such as DiffDock (Corso *et al*., 2023) and DiffDock-PP (Ketata *et al*., 2023). Finally, DiffPack (Zhang *et al*., 2024), a recent protein side-chain packing model, utilizes a confidence model. Thus, scoring models are still very useful supplemental tools in modern structural bioinformatics.

### 1.4. The specific challenges of antibody-antigen binding physics

Antibodies remain an important area of study in the development of biologics for the treatment of a variety of diseases and deep learning techniques for the design and development of antibody binders to specific epitopes is currently a very active area of deep learning research (Kim *et al*., 2023) with generative pipelines often using some form of scoring function. Furthermore, antibody interface scoring is particularly difficult due to the immense amount of structural variation in heavy chain H3 loops with little human-detectable pattern (Marks and Deane, 2017). Thus, this challenging problem may better be assisted by a specialized scoring function to help assess model generations.

## 2. Results

### 2.1. An Euclidean graph neural network architecture utilizing equivariant attention operations leveraging protein natural language model embeddings

EuDockScore, our latest scoring function, uses the equivariant attention-based architecture developed in the Equiformer and EquiformerV2 models (Liao and Smidt, 2023; Liao *et al*., 2023). These architectures have achieved state-of-the-art performance in many molecular graph learning tasks, showing increased expressivity in comparison to other graph-based architectures. In these architectures, nodes are initialized with scalar and vector features of orders *l*, where the dimensionality of the feature is given by *d* = 2*l*+1, and there may be an arbitrary number of each feature scalar or vectors. Different orders of features can equivariantly, or invariantly update each other via tensor product operations first introduced in (Thomas *et al*., 2018). The Equiformer (Liao and Smidt, 2023; Liao *et al*., 2023) introduces node level attention by generation attention values from the invariant scalar node features. Relative positional information is encoded by using the tensor product between node features and the spherical harmonics (Ledesma and Mewes, 2020) of relative positional vectors. Two major modifications were made to the EquiformerV2 architecture for this model. We moved from an all-atom representation to a residue-level representation for computational efficiency. Secondly, we input complexes rather than single polypeptide chains, and finally, we provide scalar NLP embeddings to the model instead of just raw coordinates and atom/residue identities in the form of a one-hot encoding. Please consult the EquiformerV2 (Liao *et al*., 2023) manuscript for further details.

### 2.2. An interacting protein-pair natural language model to supplement structure-based deep learning models

Large language models that implement the transformer architecture (Vaswani *et al*., 2017) have revolutionized natural language processing and immediately were put to use in learning protein embeddings using the plethora of available sequencing data. In particular, the models introduced by Elnaggar et al. (Elnaggar *et al*., 2022) and more recently Meta’s ESM2 (Lin *et al*., 2022) have been impactful to the field of protein bioinformatics. Single sequence protein folding models illustrate the power of these models’ sequence embeddings providing evidence that NLP models trained on sequence data are implicitly learning about protein folding physics, for example, ESMFold (Lin *et al*., 2022). Furthermore, protein contact maps have been shown to be recoverable to some extend from NLP embeddings (Wang *et al*., 2022). Thus, NLP models are learning important protein physical ideas that can be applied to the task of scoring input structures.

Similarly to DeepRank-GNN-esm (Xu and Bonvin, 2024), another recent scoring model, we sought to utilize NLP embeddings to supplement structure-based graph-based neural network models. We started by taking DistilProtBert (Geffen *et al*., 2022), a distilled version of the original ProtBert model (Elnaggar *et al*., 2022). We took non-redundant interacting protein sequence pairs from BioGRID (Oughtred *et al*., 2020), concatenated them together with a separating token, and fine-tuned DistilProtBert to learn about protein-protein interaction physics. These NLP embeddings are then used as input scalar features.

### 2.3. EuDockScore outperforms state-of-the-art models on the challenging CAPRI score set for protein-protein docking

As in our previous work GDockScore (McFee and Kim, 2023), we first pre-trained the model using docking decoys generated from solved structures in the PDB. We then used the refined decoys generated from BM5 (Vreven *et al*., 2015) to train the original DeepRank model (Renaud *et al*., 2021) to fine-tune the model to better understand backbone flexibility in docking and interface formation. The CAPRI score set (Lensink and Wodak, 2014) still remains the gold-standard challenging test set for assessing protein-protein docking scoring functions and we again use it as our primary benchmarking set. We chose to benchmark EuDockScore against our previous model GDockScore, and the latest iteration of the DeepRank (Renaud *et al*., 2021; Réau *et al*., 2022) scoring model recently published in the literature known as DeepRank-GNN-esm (Xu and Bonvin, 2024) referred to solely as DeepRank for the remainder of this manuscript for brevity. Furthermore, we also compare our deep learning models to the more traditional machine learning scoring function, iScore (Geng *et al*., 2019). To compare the models, we used the metrics of area under the curve (AUC) of the receiver operating characteristic (ROC) known as the AUCROC as well as compared precision-recall curves (Fig. 1a,b). Additionally, we provide the maximum Matthews correlation coefficient (MCC) and F1 score across all thresholds. We found that in comparison to DeepRank and iScore, EuDockScore and GDockScore had significantly better AUCROCs (*p <* 2.2*e* − 16) for both comparisons indicating EuDockScore more effectively can select out true positives without undesirable false positives. In comparison to each other, EuDockScore and GDockScore have similar AUCROCs. This indicates that there is a threshold where we get more true positives and less false positives in comparison to GDockScore. However, when applying statistical testing, the difference in AUCROCs is not statistically significant (*p* = 0.2468). EuDockScore has the highest MCC and F1 score across all thresholds in comparison to all models. However, we still use this as evidence of the improved discriminatory power of EuDockScore in comparison as although areas under curves may be similar the distribution of the “density” of this area may be different and important such as in the case of ROCs. The MCCs for GDockScore, EuDockScore, DeepRank, and iScore are 0.458. 0.538, 0.166, 0.279. The F1 scores for GDockScore, EuDockScore, DeepRank, and iScore are 0.528, 0.588, 0.270, 0.369. We further sought to investigate how the scores of near-native and incorrect data points where distributed by EuDockScore using a violin plot (Fig. 1c). We can see from the violin plot that EuDockScore very confidently applies high scores to near-native decoys but is less confident (indicated by a larger spread of the distribution of scores) in incorrect decoys. Logically, this is because binding is highly specific and inherently will have very precise features in comparison to the infinite number of incorrect interfaces.

**Fig. 1:**
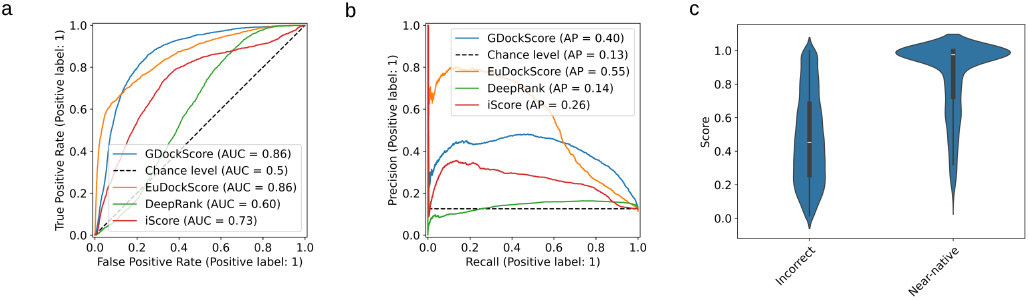
(a) ROC curves comparing EuDockScore, GDockScore, DeepRank, and iScore. (b) Precision-recall curves of EuDockScore, GDockScore, Deeprank and iScore. (c) Violin plot comparing the distribution of scores produced by EuDockScore for the CAPRI score set for near-native and incorrect decoys.

Biologists only want to experimentally validate or study a handful of structure models they are confident are near-native, so a scoring model’s ability to place near-native decoys in the top *N* decoys is important. Typically small values of *N* are selected such as 5, or 10 but we choose to graphically display the number of hits in the top *N* selected structures (Fig. 2) to be more comprehensive. In Fig. 2 below we can see that EuDockScore may do a better job on some structures (more near-native structures in a lower *N*) such as T30 and T40 but does worse on others. Ensembling improved overall ranking ability (Fig. 2) suggesting that multiple deep learning models contribute useful information to ranking proposed multimeric structural models.

**Fig. 2:**
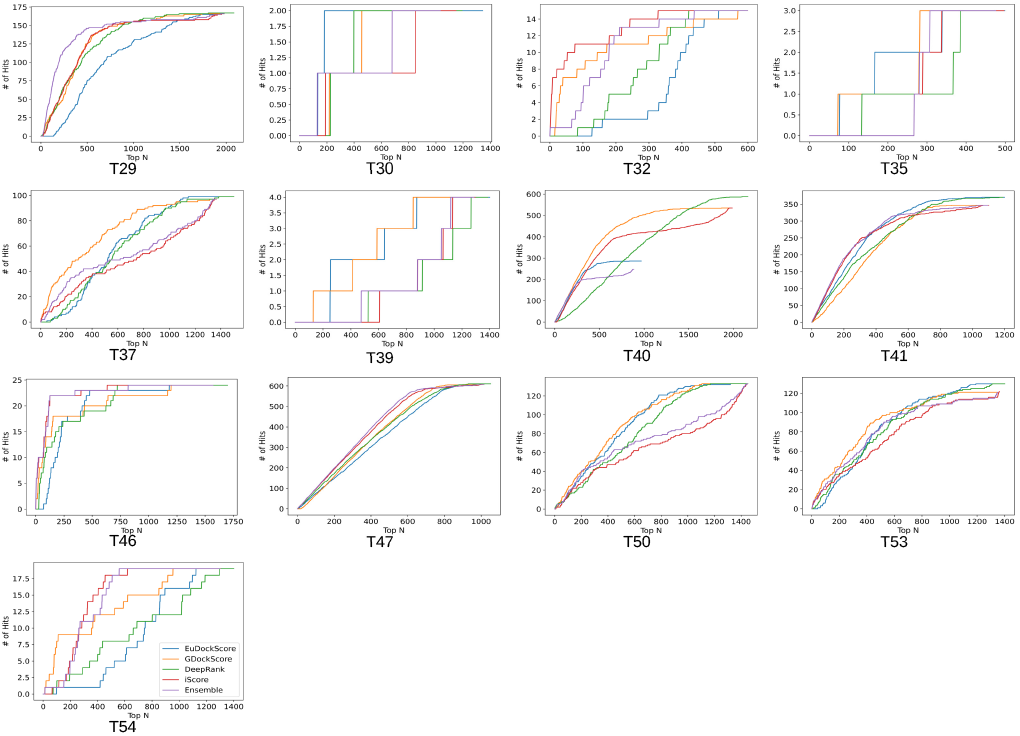
Number of hits in selected models vs. top *N* selected curves for EuDockScore, GDockScore, DeepRank, iScore, and an ensemble average of all four models. Here a “hit” is defined as a near-native structure with a CAPRI quality of acceptable or better.

### 2.4. EuDockScore performs well on CASP15 models generated by a variety of methods

To further test the capabilities of our model, we tested dimer models from CASP15 which included traditional modeling techniques and generative modeling including systems based on AlphaFold. We found that EuDockScore can differentiate high-quality vs. low-quality models for this set. We achieve an AUCROC of 0.81, an AP of 0.92 (Fig. 3 a,b), an MCC of 0.423, and an F1 of 0.868. Here we define a high-quality model as having a DockQ score >= 0.23. DockQ (Basu and Wallner, 2016) was selected as it combines many traditional model interface quality metrics into a single comprehensive metric. Furthermore, we find that our scores do correlate with DockQ scores and this correlation is statistically significant (Fig. 3c).

**Fig. 3:**
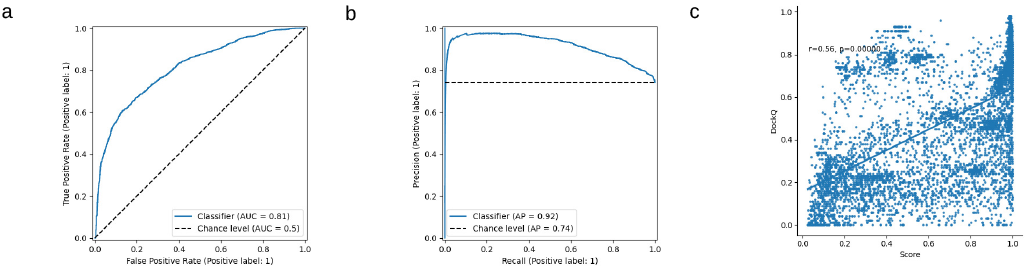
(a) ROC curve for EuDockScore for dimers in the CASP15 competition. (b) Precision-recall for dimers in the CASP15 competition. (c) Scatter plot between EuDockScore scores and DockQ values. The r value and associated p-value are indicated on the plot.

### 2.5. EuDockScore-Ab: a novel antibody-antigen complex scoring model trained on antibody-antigen docks

Next, we sought to extend EuDockScore beyond standard protein-protein docking, into the realm of assessing antibody-antigen interfaces, a task still proving to be computationally very challenging. We fine-tuned the previous EuDockScore model to produce EuDockScore-Ab, an antibody-specific model on a dataset of docked antigen-antibody structures (see Methods). Using a hold-out, non-redundant test set we see that EuDockScore-Ab can differentiate near-native from incorrect docks in unseen data (Fig. 4a,b). It is important to note our model is specific to scoring heavy chain and antigen complexes, as we found no benefit of training and using an additional light chain-specific model. Furthermore, we found that the model can extrapolate to AlphaFold-Multimer generations but does not surpass the performance of AlphaFold-Multimer in our AUCROC or AP (Fig. 5a,b). This is likely due to the end-to-end nature of the AlphaFold-Multimer training protocol and overall model size and expressiveness.

**Fig. 4:**
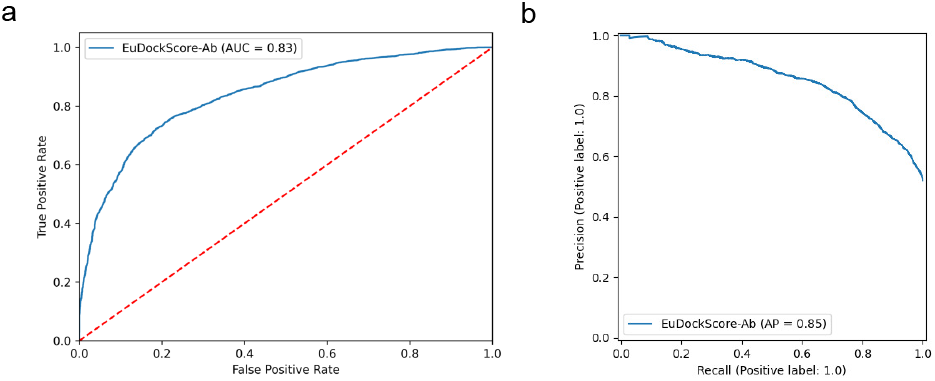
(a) ROC curve for EuDockScore-Ab on hold-out, non-redundant antibody docks. (b) Precision-recall curve for EuDockScore-Ab on hold-out, non-redundant antibody docks.

**Fig. 5:**
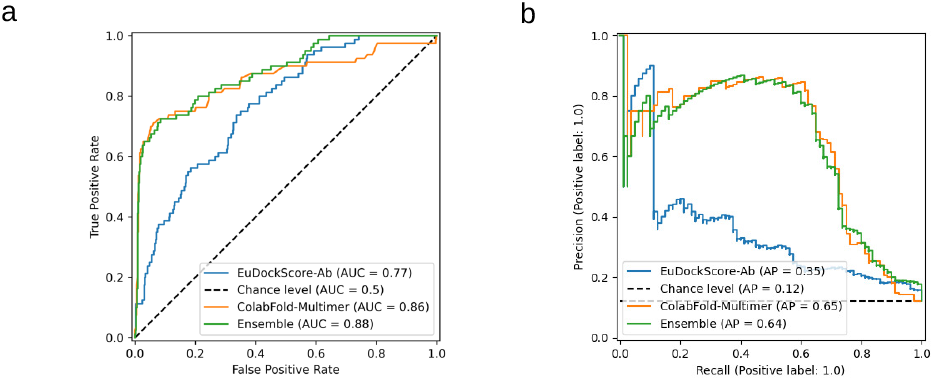
(a) ROC curves comparing EuDockScore-Ab, ColabFold-Multimer (AFM ranking confidence (0.8*ipTM + 0.2*pTM)), and an ensemble of both scoring metrics (b) Precision-recall curves comparing EuDockScore-Ab, ColabFold-Multimer (AFM ranking confidence), and an ensemble of both scoring metrics.

### 2.6. EuDockScore-AFM: attempting to distil the scoring abilities of AFM type architectures with ground-truth correction

Since AlphaFold was previously used to rank monomeric structures (Roney and Ovchinnikov, 2022), GDockScore was successful in filtering AFM outputs (Tomezsko *et al*., 2024) we next decided to distill AFM’s scoring abilities into an independent model. Instead of engineering AlphaFold2 to output single chain like scores by utilizing decoys as templates such as AF2Rank (Roney and Ovchinnikov, 2022), we propose learning a new model.

Instead of simply training to distill AFM, the new training target is the sum of the AFM ranking confidence of the model and the DockQ against the native (true) structure. This combined metric allows our model to “correct” regions of the AFM energy function using ground truth structural data. When taking this new model, hereby known as EuDockScore-AFM, and testing it on a hold-out test set of non-redundant antibody structures generated by AFSample. It is important to note that we only use the heavy chain and antigen as inputs to our model as before. Although our model does possess ranking ability according to AUCROC and AP of the precision-recall curve (Fig. 6a,b), we do not outperform the stock AFM scoring metric or pDockQ2 (Zhu *et al*., 2023), a recent scoring metric constructed from AFM quality predictions. Pairwise comparisons of all combinations of models result in statistically significant differences in AUCROC with *p* ≤ 2.2*e* − 16 for all comparisons. When we look at EuDockScore’s ranking ability for individual DockQ quality classifications, there is a trend that higher quality structures have higher EuDockScore values (Fig. 6c). For AFSample, pDockQ2, EuDockScore, and the ensemble, the F1 scores are 0.631, 0.625, 0.324, and 0.587. The maximum MCCs achieved in the same order are 0.642, 0.645, 0.268, and 0.594.

**Fig. 6:**
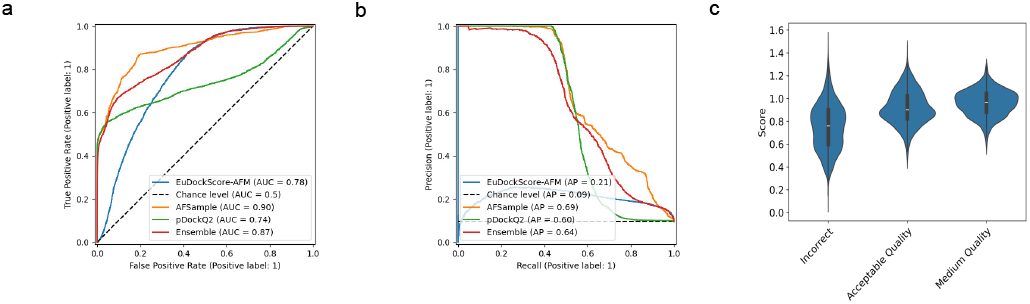
(a) ROC curves comparing EuDockScore-AFM, AFSample (AFM ranking confidence (0.8*ipTM + 0.2*pTM)), pDockQ2, and an ensemble of both scoring metrics (b) Precision-recall curves comparing EuDockScore-Ab, AFSample (AFM ranking confidence), and an ensemble of both scoring metrics.

More interestingly, when we actually compare the ranking abilities of each model for AFSample outputs, as in Figure 3. We find that there are many cases in which EuDockScore has superior ranking ability such as PDB IDs 1MLC, 5WK3, and 6A77 (Figs. 7,8). Ensembling all scoring systems also seems to help in many cases. This suggests that our model is useful in the context of helping re-rank model outputs from generative models such as AFM which has been shown in additional studies (Tomezsko *et al*., 2024). Therefore, although current scoring models do not outright replace the quality metrics of generative models such as AFM, they do provide useful information for additional filtering of structure candidate models.

**Fig. 7:**
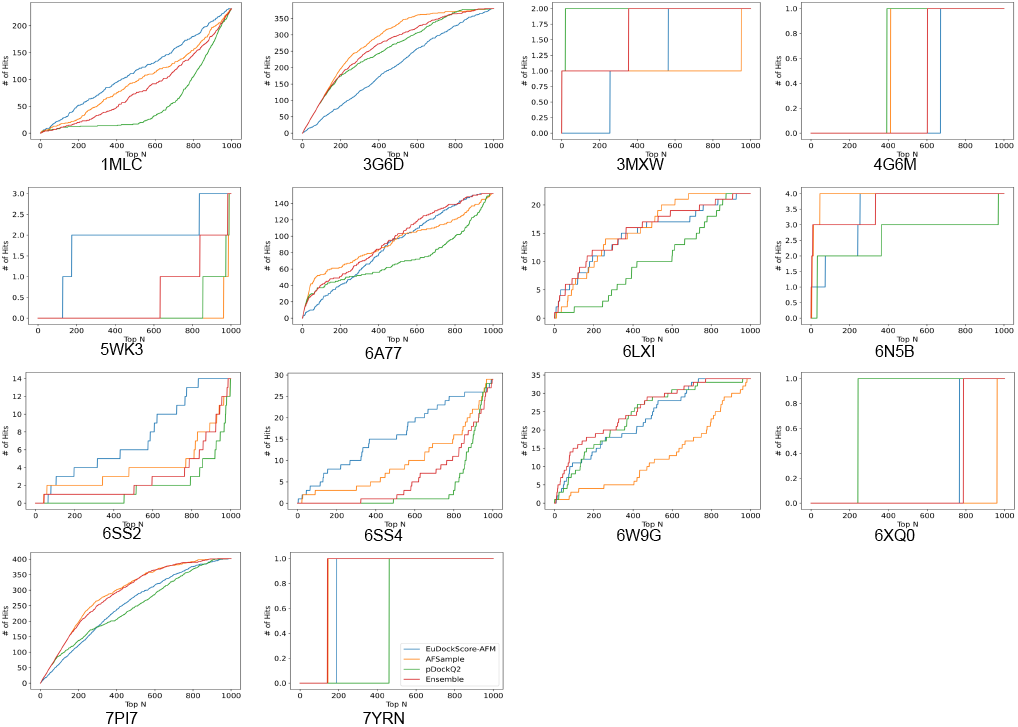
Number of hits in selected top *N* models vs. top *N* selected models for EuDockScore-AFM, AFSample, pDockQ2, and an ensemble average of all four models. Here an acceptable model or “hit” has DockQ ≥ 0.23.

**Fig. 8:**
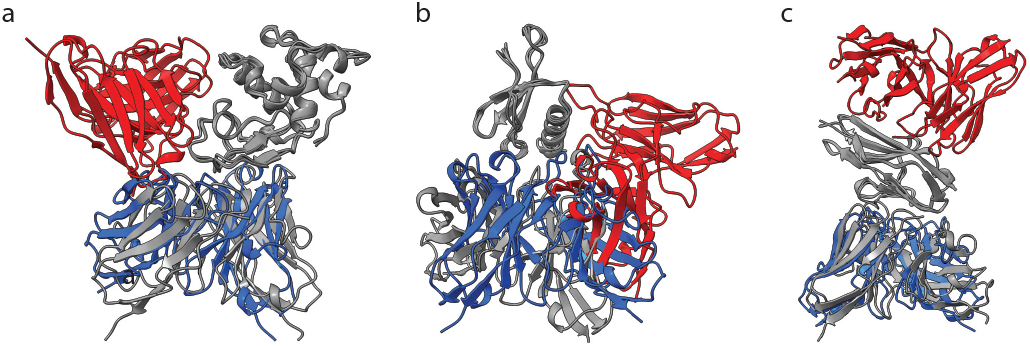
Example structures of (A) 1MLC (B) 5WK3 and (C) 6A77. Structures are superimposed based on the antigen structure. Here, grey indicates the experimentally defined PDB structure. Blue indicates structures with good EuDockScore-AFM and DockQ, but poor AFM ranking confidence. Red indicates structures with poor EuDockScore-AFM and DockQ, but good AFM ranking confidence.

## 3. Discussion

Here we have presented several EuDockScore scoring functions that can be used to accurately score protein-protein docks as well as antibody-antigen complexes. We iterate upon our previous work (McFee and Kim, 2023) by using modern, increasingly expressive Euclidean graph networks that can operate on raw coordinates (Liao and Smidt, 2023; Liao *et al*., 2023) and supplemented the inputs with NLP embeddings from a BERT-style model (Geffen *et al*., 2022).

We continue to achieve state-of-the-art performance on the CAPRI score set (Lensink and Wodak, 2014) and find that ensembling with other models such as DeepRank-GNN-esm (Xu and Bonvin, 2024), and iScore improves overall discriminative power suggesting the importance of using multiple deep learning models to assess the quality of generative modeling predictions. Our base model also generalizes well to models submitted to CASP15 further supporting the generalizability of our scoring system.

EuDockScore-Ab is a heavy chain-antigen specific scoring model trained on re-docked and relaxed real antibody complexes. EuDockScore-Ab can differentiate near-native (acceptable or better by traditional CAPRI metrics) from non-native antibody complexes. Additionally, EuDockScore-AFM a model trained specifically on AFM antibody-antigen complex predictions shows the improved ability to select out near-native structures in large pools of candidates that the standard AFM scoring metrics provided by ColabFold and AFSample showing the continued applicability of scoring function development. Although our scoring models prove useful, it is clear a significant degree of work must be done in the antibody space to achieve a state-of-the-art replacement for AlphaFold-Multimer metrics.

The work presented in this manuscript can be extended in many ways. For example, AFM models could be generated for the entire PDB to create an AFM-specific scoring function that isn’t limited to antibodies. However, this will be a computationally intensive task. Recent literature has also suggested that transformers can be applied directly to atomistic data while achieving approximate equivariance. Transformers are significantly more scalable than current equivariant networks and may allow for improved learning.

## 4. Methods

### 4.1. Generation of an antibody-antigen structure dataset using docking

Due to our previous success with GDockScore and the pre-training regime we established (McFee and Kim, 2023). We decided to replicate this training scheme for an antibody-antigen complex dataset. We used the AbDb (Ferdous and Martin, 2018), an antibody database extracted from the PDB with comprehensive processing and redundancy analysis amounting to several thousand non-redundant antibodies. Although we simply fine-tuned the previous pre-trained EuDockScore on the GDockScore and DeepRank decoys, we sought to increase antibody-specific training data complexity with re-docking and structural relaxation. To generate our dataset from AbDb, we performed local docking with the RosettaDock protocol (Gray *et al*., 2003).

We used the CAPRI standard assessment for docking quality as specified in Mendez et al. (Méndez *et al*., 2003). We assign non-native decoys as label 0 and acceptable or better decoys as label 1. We generated roughly 50000 docking decoys across the AbDb with a 70%, 15%, and 15% split to training, validation, and test sets. Redundancy information is provided by AbDb (Ferdous and Martin, 2018) and we ensured that no redundant structures were shared amongst the splits to ensure unbiased performance reporting. To further improve the quality of the interfaces and introduce back-bone flexibility into our docking models we ran the RosettaRelax protocol (Tyka *et al*., 2011) on the docking decoys to improve structure quality.

For this dataset, to ensure no redundancy between train, validation, and test sets we used the redundancy information provided by the AbDb (Ferdous and Martin, 2018) to split the AbDb into train/validation/test splits.

### 4.2. Generation of an antibody-antigen complex dataset using AFM and AFSample

To generate AFM predictions of Ab complexes, we used the protocol above, but instead generated models using the localcolabfold implementation of ColabFold (Mirdita *et al*., 2022) with templates enabled and all other options set to the default settings. Again, we split the data to avoid redundancies by clustering antigens by 30% sequence similarity with MMSeqs2 (Steinegger and Söding, 2017) and making sure there were no cluster overlaps between PDBs used in model training and validation and those used for testing. To generate a test set with a large enough number of models for meaningful re-rankings, we used AFSample (Wallner, 2023) to generate a thousand models for each PDB assigned to the test set.

### 4.3. Details of the input representation

The inputs to EuDockScore are all the raw 3D coordinates of each amino acid residue, as well as one hot encoding of amino acid identity, a binary indicator if the amino acid is within 16 angstroms of an amino acid on the other protein, as well as associated amino acid embeddings from our fine-tuned DistilProtBert model. The scalar and vector input features are projected down using equivariant operations and used to initialize irreducible node features used in the Equiformer architectures (Liao and Smidt, 2023; Liao *et al*., 2023).

### 4.4. Dataset redundancy analysis for BioGRID data

For the BioGRID dataset used to train the NLP model presented in this paper, we used the MCODE algorithm (Bader and Hogue, 2003) to cluster BioGRID and then split the data such that there was no cluster overlap in the interactions in the training and validation set.

### 4.5. Model implementation, hyperparameter tuning, and training details

Hyperparameter tuning was performed by adjusting the number of spherical channels ie. the number of types of each order l feature for each node in the EquiformerV2 (Liao *et al*., 2023) architecture as well as adjusting the maximum channel order of the type-l features of the model. We would that lmax = 2 and 16 spherical channels. Three EquiformerV2 layers are used in the model. The optimal hyperparameter configuration is set to the default configuration in the model implementation. Furthermore, we trained the model using the Adam optimizer (Kingma and Ba, 2015). The cross-entropy loss was used for the loss function during training with binary labels as specified in (McFee and Kim, 2023) and the mean-squared-error loss for training on continuous targets as in EuDockScore-AFM.

### 4.6. Definition of average precision

The average precision reported is from the implementation provided by the scikit-learn package (Pedregosa *et al*., 2011).

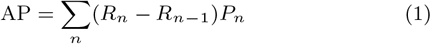

Here AP represents average precision, the summation is over the n thresholds, *R*_*n*_ is the recall at the given threshold, *R*_*n*−1_ at the previous threshold, and the weighting term *P*_*n*_ is current precision.

### 4.7. Data analysis and statistical Testing

The ROCs, precision-recall curves, Matthews correlation coefficients, and F1 scores in this paper were computed using scikit-learn (Pedregosa *et al*., 2011). The AUC of the ROC curves was compared using the DeLong test (DeLong *et al*., 1988) provided by the pROC R (R Core Team, 2021) package (Robin *et al*., 2011). The Bonferonni correction was used for multiple comparisons.

## 6. Author contributions statement

The model architecture was conceptualized by MM and PMK. Model implementation was conducted by MM. Data generation and data analysis were performed by MM and JK. Writing was performed by MM, JK, and PMK. Supervision and funding acquisition was by PMK.

## 7. Acknowledgments

We would like to thank the Digital Research Alliance of Canada for providing computing resources. CIHR Project Grants PJT-159750 and PJT-166008 funded this work.

## 8. Data and code availability

The data set introduced in this paper is available at http://gdockscore.ccbr.proteinsolver.org including the language model weights/data splits. The CAPRI score set files used in this paper are available at https://data.sbgrid.org/dataset/684/. The DeepRank data set is available at https://data.sbgrid.org/dataset/843/. The original AbDb data is available at http://www.abybank.org/abdb/. The CASP15 data, including DockQ values, was download from https://predictioncenter.org/download_area/. The code is available at https://gitlab.com/mcfeemat/eudockscore.

